# Seasonal variability in carbon:^234^thorium ratios of suspended and sinking particles in coastal Antarctic waters: Field data and modeling synthesis

**DOI:** 10.1101/2022.01.07.475468

**Authors:** Michael R. Stukel, Oscar M. E. Schofield, Hugh W. Ducklow

## Abstract

^238^U-^234^Th disequilibrium is a powerful tool for investigating particle cycling and carbon export associated with the ocean’s biological carbon pump. However, the interpretation of this method is complicated by multiple processes that can modify carbon:thorium ratios over small spatial scales. We investigated seasonal variability in the thorium and carbon cycles at a coastal site in the Western Antarctic Peninsula. Throughout the ice-free summer season, we quantified carbon and ^234^Th vertical flux, total water column ^234^Th, particulate ^234^Th, and the C:^234^Th ratios of sinking material and bulk suspended material. Simultaneous identification and separation of fecal pellets from sinking material showed that fecal pellets (primarily from krill) contributed 56% of carbon flux and that as a result of lower C:^234^Th ratios than suspended particles, these fecal pellets were primary drivers of variability in the C:^234^Th ratios of sinking material. Bulk suspended particles had highly variable C:^234^Th ratios and were consistently elevated in the euphotic zone relative to deeper waters. The fraction of ^234^Th adsorbed onto particles was positively correlated with chlorophyll and particulate organic carbon (POC) concentrations. The C:^234^Th ratios of suspended particles were positively correlated with POC, although during the spring diatom bloom C:^234^Th ratios were lower than would have been predicted based on POC concentrations alone. We hypothesize that diatom production of transparent exopolymers may have led to enhanced rates of thorium adsorption during the bloom, thus decreasing the C:^234^Th ratios. We used a Bayesian model selection approach to develop and parameterize mechanistic models to simulate thorium sorption dynamics. The best model incorporated one slowly-sinking POC pool and rapidly-sinking fecal pellets, with second-order sorption kinetics. The model accurately simulated temporal patterns in the C:^234^Th ratios of sinking and suspended particles and the fraction of ^234^Th adsorbed to particles. However, it slightly over-estimated C:^234^Th ratios during the spring (diatom-dominated) bloom and underestimated C:^234^Th ratios during the fall (mixed-assemblage) bloom. Optimized model parameters for thorium sorption and desorption were 0.0047 ± 0.0002 m^3^ mmol C^-1^ d^-1^ and 0.017 ± 0.008 d^-1^, respectively. Our results highlight the important role that specific taxa can play in modifying the C:^234^Th ratio of sinking and suspended particles and provide guidance for future studies that use ^234^Th measurements to investigate the functional relationships driving the efficiency of the biological pump.

**HIGHLIGHTS:** Investigated thorium and carbon cycling over full ice-free season

C:^234^Th ratios of sinking particles were controlled by low C:^234^Th of fecal pellets

C:^234^Th ratios of suspended particles were correlated with chlorophyll and POC

Diatom abundance may have led to high particulate thorium during spring bloom

Second-order thorium sorption kinetics model accurately simulates C:^234^Th ratios

## 1. INTRODUCTION

Photosynthesis in the surface ocean decreases the partial pressure of CO_2_ in the oceanic mixed layer, leading to CO_2_ uptake from the atmosphere (Falkowski et al., 1998; Raven and Falkowski, 1999). However, as a result of the short life spans of pelagic primary producers, most CO_2_ taken up is respired back into the surface ocean. Long-term carbon sequestration is dependent on processes that transport organic matter from the surface to the deep ocean and are collectively referred to as the biological carbon pump (BCP) (Boyd *et al*., 2019; Ducklow *et al*., 2001; Volk and Hoffert, 1985). The BCP is often driven by the gravitational settling of organic particles and aggregates (Boyd *et al*., 2019; Buesseler and Boyd, 2009), although recent evidence highlights the spatiotemporally variable importance of organic matter subduction and active transport by vertical migrants (Archibald *et al*., 2019; Bianchi *et al*., 2013; Omand *et al*., 2015; Stukel *et al*., 2018). Current estimates of the global magnitude of the BCP range from 5–13 Pg C yr^-1^ (Dunne *et al*., 2005; Henson *et al*., 2011; Laws *et al*., 2011; Siegel *et al*., 2014). Our ability to quantify interannual variability in the BCP, or predict its response to a changing climate, is hampered by methodological difficulties that currently inhibit large-scale monitoring of the BCP (McDonnell *et al*., 2015).

Two commonly used approaches for quantifying sinking particle flux are sediment traps and radionuclide disequilibria pairs. Sediment traps are conceptually simple tools that involve the deployment and recovery of instruments capable of collecting sinking particles (Buesseler *et al*., 2007; Honjo *et al*., 2008; Martin *et al*., 1987). Sediment traps are commonly deployed in either long-term moored, surface-tethered, or neutrally-buoyant configurations. Long-term moored traps have been extensively used for monitoring of particle flux near the seafloor (Honjo *et al*., 2008), however when deployed at shallow depths to quantify particle flux entering the mesopelagic zone they are prone to substantial biases (Buesseler *et al*., 2010). Surface-tethered and neutrally-buoyant traps may provide better results in shallow water (Baker *et al*., 2020), but require substantial ship-time investment for short-term deployments.

Radionuclide disequilibrium pairs (e.g., ^238^U-^234^Th) provide an alternate approach for quantifying flux that is well-suited for the survey sampling plans of many oceanographic programs (Le Moigne et al., 2013; Van der Loeff et al., 2006; Verdeny et al., 2009; Waples et al., 2006). The parent nuclide (^238^U) is a conservative element that decays into a shorter-lived, particle-reactive nuclide (^234^Th). When the daughter particle is scavenged onto particles and removed from the surface ocean by particle sinking, it creates a disequilibrium between parent and daughter that reflects the magnitude of particle flux that has occurred over the decay lifetime of the daughter particle (Buesseler *et al*., 1992; Coale and Bruland, 1985; Santschi *et al*., 2006). The use of simple steady-state models or more complex non-steady state models that may incorporate physical mixing and advection then allow vertical profiles of disequilibrium to be used to quantify radionuclide flux (Dunne and Murray, 1999; Resplandy et al., 2012; Savoye et al., 2006). Estimations of carbon or nitrogen flux then require measurements of the C:radionuclide or N:radionuclide ratio of sinking particles (Buesseler *et al*., 2006; Puigcorbé *et al*., 2020).

C:^234^Th ratios can be highly variable even over relatively small spatial scales (Buesseler *et al*., 2006; Hung and Gong, 2010; Stukel *et al*., 2019). This variability is driven by complex chemical, ecological, and physical interactions. Thorium adsorbs onto particle surfaces, suggesting that C:^234^Th ratios should increase with increasing particle size due to the decreased surface area:volume ratio of large particles.

However, likely as a result of the heterogeneity of particle types in the ocean and multitude of processes that reshape the particle size spectrum, simple relationships between particle size and C:^234^Th ratios have proved elusive (Buesseler et al., 2006; Burd et al., 2007; Hung and Gong, 2010; Passow et al., 2006). ^234^Th scavenging is also affected by the presence of acid polysaccharides that have multiple strong sorption sites for ^234^Th (Guo et al., 2002; Passow et al., 2006; Quigley et al., 2002; Zhang et al., 2008) and by the relative partitioning between sinking particles and suspended particles and colloids (Murphy *et al*., 1999). C:^234^Th ratios in large size fractions or sediment trap samples can also be affected by the presence of mesozooplankton, which tend to have very high C:^234^Th ratios (Coale, 1990; Dunne et al., 2000; Passow et al., 2006; Stukel et al., 2016; Stukel et al., 2019) or mesozooplankton fecal pellets, which might have low C:^234^Th ratios if zooplankton assimilate carbon but not ^234^Th (Rodriguez y Baena *et al*., 2007; Stukel *et al*., 2019).

Because of the multiple factors that complicate *a priori* prediction of the C:^234^Th ratio of sinking particles, it is ideal to empirically determine these ratios for sinking particles at every location and time for which particle flux estimates are desired. However, typical oceanographic survey designs often do not permit such sampling plans, leading researchers to interpolate coarse C:^234^Th ratio measurements (or no contemporaneous measurements at all) to finer resolution water column ^234^Th sample locations (Ducklow et al., 2018; Estapa et al., 2015; Puigcorbé et al., 2017; van der Loeff et al., 2011). Clearly additional information relating C:^234^Th ratios to contemporaneous chemical, physical, and biological parameters is necessary to enable more accurate estimation of carbon (or other element) flux at times when this parameter cannot be measured.

One potentially fruitful approach for estimating spatiotemporal variability in C:^234^Th ratios at multiple scales is the development of mechanistic, coupled ^234^Th sorption models (Resplandy *et al*., 2012). Such models have been used for diverse purposes including quantifying the impacts of physical circulation on ^234^Th export estimates in the Equatorial Pacific (Dunne and Murray, 1999), investigating the impacts of aggregation on ^234^Th flux (Burd *et al*., 2007), illustrating mesoscale variability in ^234^Th activity (Resplandy *et al*., 2012), simulating deepwater sorption processes (Lerner *et al*., 2016), and quantifying the impact of particle sinking speeds on vertical distributions of C:^234^Th ratios (Stukel and Kelly, 2019). Despite their increasingly frequent use, there is substantial uncertainty about what model compartments (i.e., state variables) are necessary to accurately simulate the system. Furthermore even the basic functional forms for modeling thorium sorption remain in question. For instance, Lerner *et al*. (2016) assumed sorption was driven by first-order kinetics (i.e., sorption rates are proportional to dissolved ^234^Th) but concluded that thorium sorption coefficients were depth-dependent, while Replandy *et al*. (2012) and Stukel and Kelly (2019) applied second order kinetics (i.e., sorption rates are proportional to the product of dissolved ^234^Th×POC) with non-varying sorption coefficients. The specific parameters associated with thorium sorption and desorption, and whether or not these parameters differ for different classes of particles, remain highly uncertain.

This study focuses on seasonal variability in thorium and carbon cycling at a coastal site in the Western Antarctic Peninsula (WAP). The coastal WAP is a heavily sea-ice influenced region in which summer diatom blooms typically support large euphausiid populations (Ducklow *et al*., 2013; Saba *et al*., 2014). Sinking particle flux in the WAP exhibits high seasonal variability and is dominated by euphausiid fecal pellets, although vertical mixing may also play an important role in the BCP (Ducklow *et* al., 2008; Gleiber *et al*., 2012; Stukel and Ducklow, 2017). We present results from weekly sampling of water column ^234^Th, water-column particulate C:^234^Th ratios, the C:^234^Th ratios of sinking particles, and sinking fecal pellet flux made in parallel with chemical and biological measurements made by the Palmer Long-Term Ecological Research (LTER) Program including chlorophyll *a* (Chl), particulate organic carbon (POC), nutrients, and primary production. We also use a Bayesian model selection approach to simulate thorium sorption and desorption processes and changing C:^234^Th ratios; to objectively evaluate models of differing complexity using either first-order or second-order sorption kinetic equations; and to estimate thorium sorption parameters.

## 2. METHODS

### 2.1. Oceanographic sampling

Sampling took place during the Palmer LTER 2012-2013 field season from October 31st (as soon as the winter sea ice cleared) until March 25^th^ (near the end of the phytoplankton growing season). Samples were primarily collected from Palmer LTER Station E, which is located 3.2 km from shore in ∼170 m water depth in the Bismarck Strait (Fig. 1). Basic physical, chemical, and biological measurements (temperature, salinity, photosynthetically active radiation, nutrients, chlorophyll, and primary productivity) were made twice weekly. Water column total ^234^Th and particulate ^234^Th were measured weekly. Moored sediment traps were typically deployed and recovered for continuous one-week deployments. However, during early and late seasons, conditions (primarily sea ice) required shorter, non-overlapping deployments.

**Fig. 1. –.**
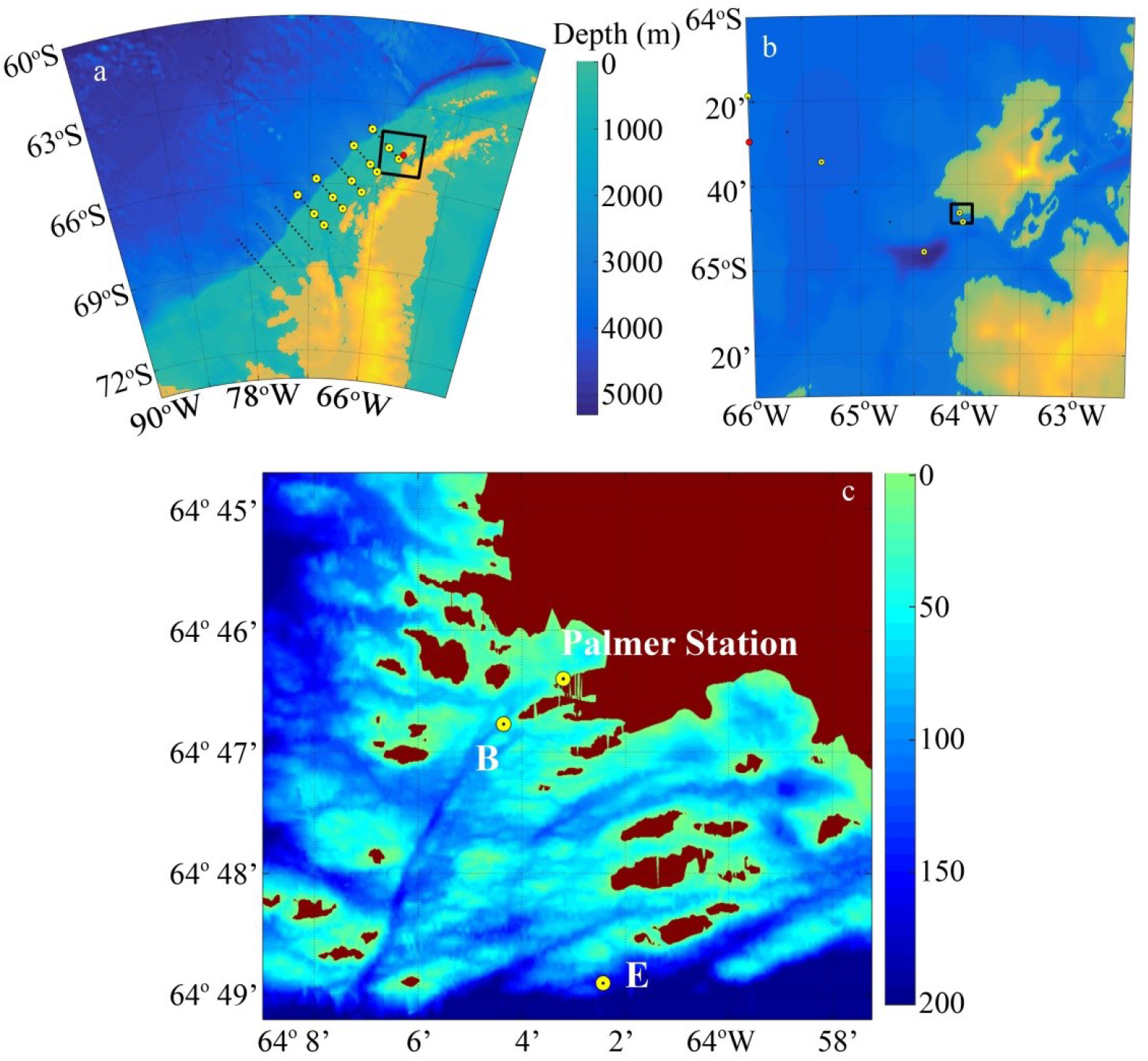
Map of the study region. a) Western Antarctic Peninsula, showing larger Palmer LTER sampling grid. b) Anvers Island region. c) Palmer LTER study region.

### 2.2 Sediment trap

We used a moored version of VERTEX-style sediment traps (Knauer *et al*., 1979; Stukel *et al*., 2015). This moored configuration was necessary, because drifting traps would rapidly have floated out of the 3.2-km radius region surrounding Palmer Station within which small boat operations are permitted. The trap array included four particle-interceptor trap (PIT) tubes (60-cm height, 7-cm internal diameter) with a baffle comprised of 13 smaller tubes with tapered ends, on a PVC cross-piece. PIT tubes were filled with saltwater brine made from 0.1-μm filtered seawater amended with 40 g L^-1^ NaCl and borate-buffered formaldehyde (final concentration 0.4%). The trap was moored 50-m from the surface in ∼170-m deep water near station E (during the middle of the season) or in ∼80-m deep water at station B (at the beginning and end of the season when conditions and circumstances did not allow deployment at E, see Fig. 1). Trap deployments during the middle of the season were of roughly one-week duration, with the trap redeployed immediately following recovery. At the beginning and end of the season sea ice increased the risk of losing the trap, so deployments were shortened to a period of 2-4 days.

After recovery, overlying less dense water was removed by gentle suction and PIT tube brine was mixed by gentle inversion. 50-150 mL aliquots were taken from each of three tubes and filtered onto GF/F filters for pigment analyses. The remainder of each of the three tubes was similarly weighed and filtered through a 200-μm filter. The filtrate was filtered through a pre-combusted QMA filter for C:^234^Th analyses of the <200-μm fraction. The material on the 200-μm filter was inspected under a Leica MZ stereomicroscope to allow removal of mesozooplankton swimmers. The remainder of the >200-μm sample was then filtered through a pre-combusted QMA filter for C:^234^Th analyses of the large size-fraction of sinking material.

The fourth PIT tube was used for fecal pellet enumeration. Samples were size-fractionated through a 200-μm filter as described above. The >200-μm size-fraction was rinsed into a gridded plastic petri dish and an average of 330 pellets per sample were enumerated under a Leica MZ 7.5 dissecting microscope at 6.3X magnification. Images were taken of random grid cells with a Nikon Digital Sight DS-Fi1c digital camera. When a visual assessment suggested that >90% of the particle volume in the sample was composed of fecal pellets, large non-fecal pellet particles were removed (though large non-fecal pellet particles were never common) and the remainder of the sample was filtered through a pre-combusted QMA filter for analysis of the C:volume and C:^234^Th ratio of fecal pellets. The <200-μm size fraction was poured into a graduated cylinder and fecal pellets were allowed to settle in the bottom of the cylinder. Liquid was then decanted through a 10-μm filter until ∼10mL remained. Remaining liquid containing sinking material was then poured into a gridded petri dish and 10-μm filter was rinsed into petri dish. Random grid cells were examined under the Leica dissecting microscope and imaged at 25X magnification. A minimum of 100 fecal pellets were imaged for each sample. Images were examined and fecal pellets were outlined using ImagePro software. Fecal pellet dimensions (length and width) were used to compute volume using the equation of a cylinder (since the vast majority of pellets were cylindrical euphausiid pellets). The empirically determined carbon density of the fecal pellets (2.15 μmol C mm^-3^) was used to estimate pellet carbon flux from microscopy images of fecal pellets.

### 2.3. Water column thorium sampling

Three types of measurements were made to quantify thorium distributions in the water column: total ^234^Th, bulk (>1-µm) particulate ^234^Th, and >50-μm ^234^Th. Total ^234^Th was measured using standard small volume techniques (Benitez-Nelson *et al*., 2001; Pike *et al*., 2005). 3-4 L samples (exact volume determined gravimetrically on land) for total ^234^Th were sampled by Go-Flo bottles from eight depths (0, 5, 10, 20, 35, 50, 65, 100 m). Samples were acidified to a pH of <2 with HNO_3_. A tracer addition of ^230^Th was added and samples were mixed vigorously. Samples were allowed to equilibrate for 4-9 hours and then adjusted to a pH of 8-9 with NH_4_OH. KMnO_4_ and MnCl_2_ were added and samples were mixed and allowed to sit for ∼12 h as Th co-precipitated with manganese oxide. Samples were then vacuum-filtered at high pressure onto QMA filters, dried, and mounted in RISO sample holders.

3-4 L particulate ^234^Th samples were similarly collected (although only at depths from 0-65 m) and weighed on land. Samples were then immediately vacuum-filtered at a pressure of 5-7” Hg onto a pre-combusted QMA filter. Filtrate was collected and 4 L was filtered through an additional pre-combusted QMA filter to serve as a blank to account for adsorption of dissolved ^234^Th or organic carbon onto filter. Filters were immediately dried in a drying oven and mounted in RISO sample holders. A gap in particulate ^234^Th sampling occurred from Jan. 17^th^ to Feb. 8^th^.

>50-μm ^234^Th particulate samples were collected by two different methods, depending on depth. At the surface, samples were collected by directly filling a 50-L carboy with surface water and gravity filtering through a 50-μm nitex mesh filter. Samples from 10 and 30 m depth were collected by a submersible pump (Proactive Industries SS Monsoon Pump) and filtered through a 50-μm nitex mesh filter on the zodiac. Volumes filtered depended on particle concentration and ranged up to 100 L. Particles were rinsed from 50-μm filters onto pre-combusted QMA filters, dried, and mounted in RISO sample holders. We note that this approach to collecting large particles and aggregates might be more likely to generate turbulence that disrupts fragile aggregates than frequently used stand-alone in situ pumps. It was the only approach that was feasible when sampling from a zodiac, however.

Samples for water column ^234^Th, particulate ^234^Th, >50-μm particulate ^234^Th, and sediment trap ^234^Th were beta counted on a RISO low-level background beta counter at Palmer Station and re-counted >6 half-lives later. Samples for water-column ^234^Th were then dissolved in HNO_3_/H_2_O_2_ solution and ^229^Th tracer was added. Samples were evaporated and reconstituted in dilute nitric acid / hydrofluoric acid. They were then analyzed by inductively-coupled plasma mass spectrometry at the Woods Hole Oceanographic Institute Analytical Lab to determine the ratio of ^229:230^Th to determine the initial yield of the ^234^Th filtration. Samples for particulate ^234^Th (water column, size-fractionated, and sediment trap) were fumed with HCl to remove inorganic carbon and combusted in a CHN analyzer to quantify particulate organic carbon (POC) and the C:^234^Th ratio.

^234^Th export was estimated from water-column ^234^Th measurements using a one-dimensional non-steady state equation that estimated vertical introduction of ^234^Th as a diffusive process:

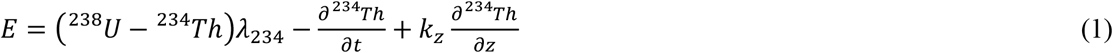

where ^238^U and ^234^Th are the activities of ^238^U and ^234^Th vertically-integrated above a reference depth (50 m here, for comparison to sediment traps), λ_234_ is the decay constant for ^234^Th, and k_z_ is a temporally-varying vertical eddy diffusivity coefficient estimated at our sampling location (Stukel *et al*., 2015). ^238^U activity was estimated from a linear relationship with salinity (Owens *et al*., 2011). ^234^Th was estimated as a two-week moving average.

### 2.4. Biological measurements

Chl *a* samples were taken at 5 depths (0, 5, 10, 20, and 65 m), filtered onto GF/F filters and measured using a fluorometer with the acidification method (Strickland and Parsons, 1972). Samples for net primary productivity (NPP) were taken from the same depths, transferred into polycarbonate bottles, and spiked with H^14^CO_3-_. Bottles were covered in mesh screening to achieve irradiance levels equal to 100%, 50%, 25%, 10%, and 0% surface irradiance and incubated in an outdoor incubator with temperature corresponding to surface seawater. These irradiance levels correspond to typical light levels at those depths during the season. However, at the height of the spring bloom, light levels were substantially lower *in situ* than in the sample bottles. Hence we corrected our NPP estimates by fitting a primary productivity model based on Moline et al. (1998): PP = P_max_×Chl×tanh(PAR/I_k_). We determined P_max_ and I_k_ from our H^14^CO_3-_ uptake incubations, combined with contemporaneous Chl *a* measurements and PAR determined by multiplying 24 hour averages of Palmer Station surface PAR by our incubation light percentages. For additional details, see Stukel et al. (2015).

### 2.5. Statistical analyses

To smooth and interpolate our unevenly sampled data fields from station E, we used the geostatistical technique of kriging (Krige, 1951). This approach was particularly important for matching total water column ^234^Th and particulate ^234^Th measurements, because these measurements were typically made 2 days apart. Linear regressions were computed using Type II geometric mean regressions (Matlab function lsqfitgm), because independent variables were never controlled. Correlations were tested using Pearson’s linear correlation (Matlab function corr).

### 2.6 Thorium sorption models

To mechanistically investigate the processes driving variability in the C:^234^Th ratio we developed simple dynamic models describing transformations of particulate organic carbon and thorium sorption and desorption. We had three primary objectives in developing these models: 1) Ascertain whether first-order kinetics or second-order kinetics more accurately described thorium sorption processes. 2) Determine what processes (e.g., fecal pellet production; differences in sorption between phytoplankton and detritus) must be included to accurately model variability in the C:^234^Th ratio. 3) Objectively parameterize key sorption rate coefficients, along with their associated uncertainty. Accurate mechanistic modeling of the complex biogeochemistry of the WAP, or indeed any marine ecosystem, is exceedingly challenging (Hood et al., 2006; Schultz et al., 2020), and our goal was not to develop a model capable of completely simulating carbon and nitrogen dynamics in the region. We therefore developed diagnostic models of particulate organic carbon. Specifically, we determined time-varying fields associated with biogeochemical transfer functions (e.g., primary productivity, remineralization, particle sinking flux) directly from our field measurements (see online supplementary appendix). These diagnostic models were able to completely model the processes driving temporal variability in POC (and phytoplankton biomass for more complex models), while exactly matching the field measurements of POC (Supp. Figs. 1 – 4). We then coupled the diagnostic POC models to mechanistic models of thorium transformations using either first-order or second-order sorption kinetics.

We used four different model structures for organic carbon (Fig. 2): particulate organic carbon only (POC); POC and krill fecal pellets (POC-FP); phytoplankton, detritus, and krill fecal pellets (Phy-Det-FP); and diatoms, flagellates, detritus, and krill fecal pellets (Dtm-Flag-Det-FP). All models included sinking (for detritus or POC) and vertical mixing (all compartments). Model 1 (POC model) also included NPP and remineralization. Model 2 (POC-FP model) added grazing by krill and the production of fecal pellets which we assume sank from the euphotic zone instantaneously. Model 3 (Phy-Det-FP model) split the POC pool into phytoplankton and other detritus (which includes heterotrophic bacteria and protists, although we assume that non-living detritus is the bulk of this pool). The Phy-Det-FP model includes NPP and mortality of phytoplankton. A portion of this mortality produces detritus, while the remainder is lost to the dissolved pool. Detritus then experiences remineralization. Model 4 (Dtm-Flag-Det-FP model) includes the same processes, except that it splits the phytoplankton compartment into diatoms and flagellates. These four carbon models were each coupled to two variants of a thorium-sorption model (first-order or second-order). First order models parameterized thorium sorption as: sorption = k_1_×^234^Th_diss_. Second-order models parameterized thorium sorption as: sorption = k_2_×POC×^234^Th_diss_. We assumed that different model carbon components could have different thorium-binding properties (e.g., for the Phy-Det-FP model we assumed that Phy and Det have different thorium sorption coefficients: k_1,Phy_ and k_1,Det_). All models also included desorption, decay, and production of dissolved ^234^Th from ^238^U. For additional description of the models, including all diagnostic equations, please see the online supplementary appendix.

**Fig. 2. –.**
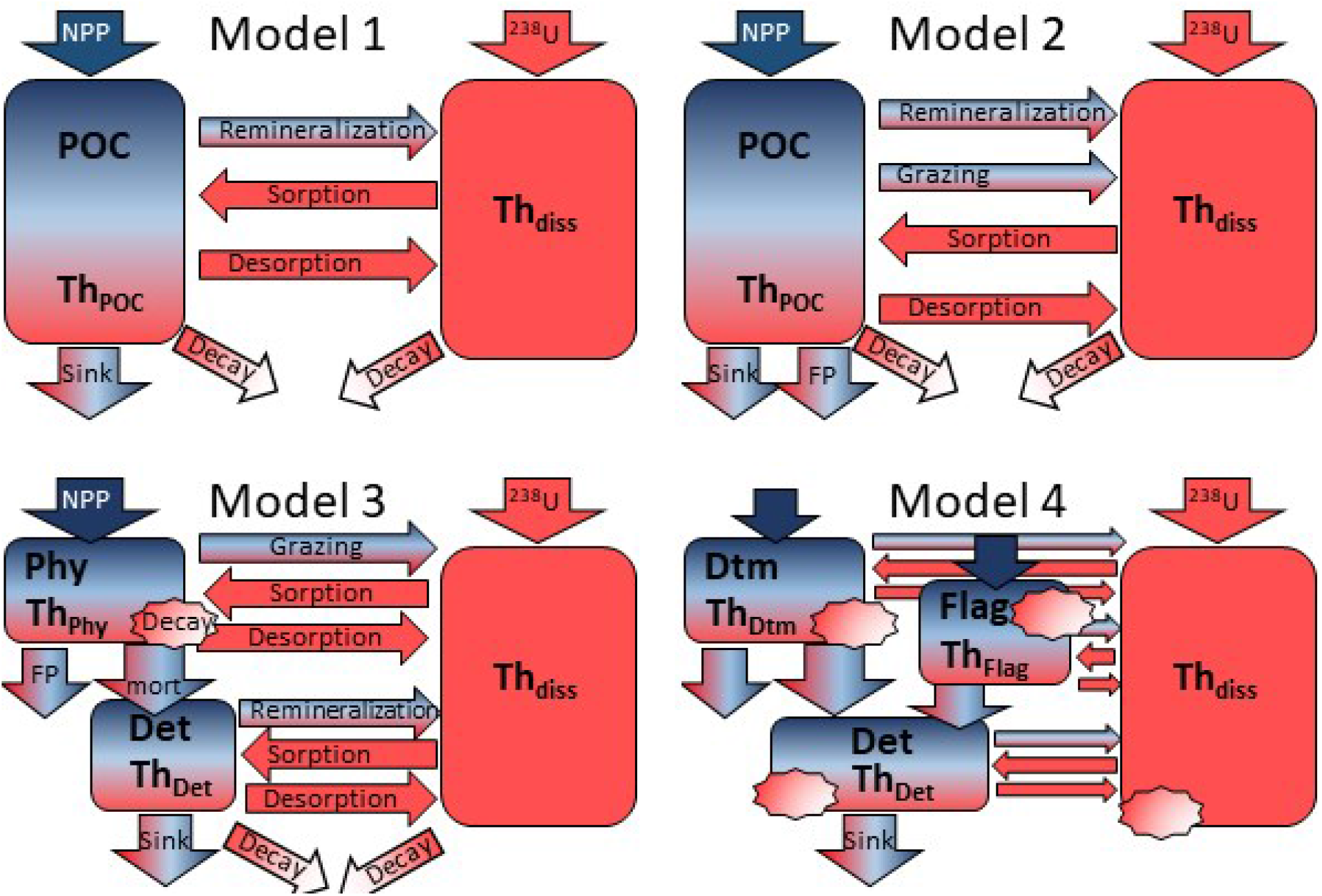
Model structure for the POC Model (Model 1), POC-FP Model (Model 2), Phy-Det-FP Model (Model 3), and Dtm-Flag-Det-FP Model (Model 4). Standing stocks (rectangles) and fluxes (arrows and callouts) are color coded based on whether they apply only to POC (blue), only to ^234^Th (red), or to both (red and blue). Note that Model 4 does not have labels associated with fluxes, but fluxes are very similar to those in Model 3 but with the phytoplankton (Phy) compartment split into diatoms (Dtm) and flagellates (Flag). For additional explanation see text and online appendix.

We ran the model in a one-dimensional configuration for the upper 65 m of the water column with 5-m thick layers and a 3-minute time step. To match our field data, we initialized the model with field data from November 10 and ran it to December 10 and then reinitialized it from Dec. 20 and ran it to March 17. We chose not to model the period from Dec. 10 – 20, because wind data showed that these dates encompassed the only period of consistent offshore winds, causing an across-shore current that invalidates the assumption of our one-dimensional model (Stukel *et al*., 2015). We note that a one-dimensional model neglects along-shore and across-shore advective processes that have the potential to impact water column standing stocks of ^234^Th and POC. However, we feel that this is justified, because: 1) a one-dimensional nitrogen-mass-balance-constrained model suggested that our sampling site could be adequately modeled as a one-dimensional system except from Dec. 10 – 20 (Stukel *et al*., 2015); 2) wind measurements and water column measurements from a nearby site did not suggest horizontal advection as a dominant term in ^234^Th or POC budgets; and 3) Markov Chain Monte Carlo parameter exploration (see below) is not computationally feasible with a three-dimensional model.

### 2.7 Bayesian model parameterization and selection

The aforementioned eight models varied substantially in their complexity. Model 1 had two unknown parameters to be fitted; Model 2 had three; Model 3 had four; and Model 4 had five. To fit these parameters to the field data, we used a Bayesian statistical framework solved using a Markov Chain Monte Carlo approach (Metropolis *et al*., 1953). Specifically, starting with an initial guess for all parameter values, we ran the model from November 10 - December 10 and from December 20 – March 17. We then computed the model misfit to all C:^234^Th data: suspended particle C:^234^Th field data was compared to the sum of organic carbon in all particulate model compartments divided by the sum of particulate ^234^Th in those same compartments. We also compared the C:^234^Th ratio of model sinking particles at 50-m depth to sediment trap C:^234^Th and for all models that included fecal pellets we also compared modeled fecal pellet C:^234^Th ratios to >200-μm sediment trap C:^234^Th and modeled small detritus sinking C:^234^Th to <200-μm sediment trap C:^234^Th. Log likelihood (ln(L)) was computed from misfit as: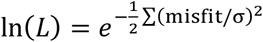, where s is measurement uncertainty.

We then proposed a new value for each parameter by drawing a random number from a normal distribution centered at the previous value for each parameter set. We re-ran the model with this new proposed parameter set and re-computed ln(L). This proposed parameter set was accepted with probability:

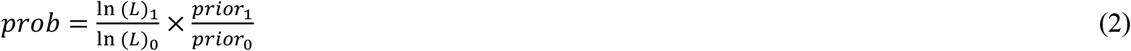

Where *prior* represents the prior density of the parameter set, and subscripts 0 represent the original parameter set, while subscripts 1 represent the proposed parameter set. For all sorption and desorption parameters, we assumed log-normal prior distributions. For first-order forward sorption coefficients we chose a prior distribution with a mean of 0.7 y^-1^ from Lerner et al. (2016). For second-order forward sorption coefficients we used a prior distribution with a mean of 0.013 m^3^ mmol C d^-1^ from Stukel et al. (2019). For the desorption coefficient we chose a prior with a mean of 2 y^-1^ (Lerner *et al*., 2016). Since the thorium egestion coefficient (Eg_Th_), which parameterizes the proportion of thorium consumed by euphausiids that is egested in their fecal pellets, must vary between zero and one, we modeled its prior with a beta distribution. We centered the beta distribution at 0.5, because of evidence that a higher proportion of thorium is egested than carbon, while carbon-based egestion efficiencies are typically ∼30% (Conover, 1966; Stukel *et al*., 2019). For all prior distributions we chose shape parameters that reflected a coefficient of variation equal to 0.5 to represent substantial uncertainty in these parameters. We ran the Markov Chain Monte Carlo simulation for a minimum of 200,000 iterations (longer if results had not stabilized) and removed the initial 10% of the iterations as a burn-in period. To objectively compare the differences between different model structures we used the deviance information criterion (DIC, Spiegelhalter *et al*., 2002). DIC accounts for varying model complexity (i.e., number of parameters) with better performing models having a lower DIC.

## 3. RESULTS

### 3.1. Conditions during the 2012-2013 field season

At the beginning of our sampling season, phytoplankton biomass was low (<2 μg Chl *a* L^-1^) and mixed-layer nitrate concentrations were high (28 μmol L^-1^, Fig. 3), typical early-season conditions at Palmer Station and other locations along the Peninsula (Kim *et al*., 2018). Phytoplankton abundance and net primary productivity remained low (and nutrients remained high) until mid-November, when a strong phytoplankton bloom formed (peak Chl *a* concentrations ∼20 μg Chl a L^-1^). High nitrate uptake rates were paired with rapid nutrient drawdown in surface layers to ∼5 μmol NO_3_^-^ L^-1^. Although strong in magnitude, the bloom was brief and terminated by early December. Although the end of the bloom coincided with peak export, export measured in the sediment traps was not sufficient to explain the rapid loss of POC from the euphotic zone. Instead, evidence (including sustained offshore favorable winds and a rapid return to high nutrient concentrations) suggested that the end of the bloom coincided with an offshore lateral advection event common in this region (Oliver *et al*., 2019). Chl, POC, and NPP remained low for approximately two months following the decay of the bloom and nutrient concentrations actually increased in the euphotic zone over this period (Fig. 3). In mid to late February, a late summer bloom began to form, although Chl concentrations never exceeded 4 μg Chl *a* L^-1^.

**Fig. 3. –.**
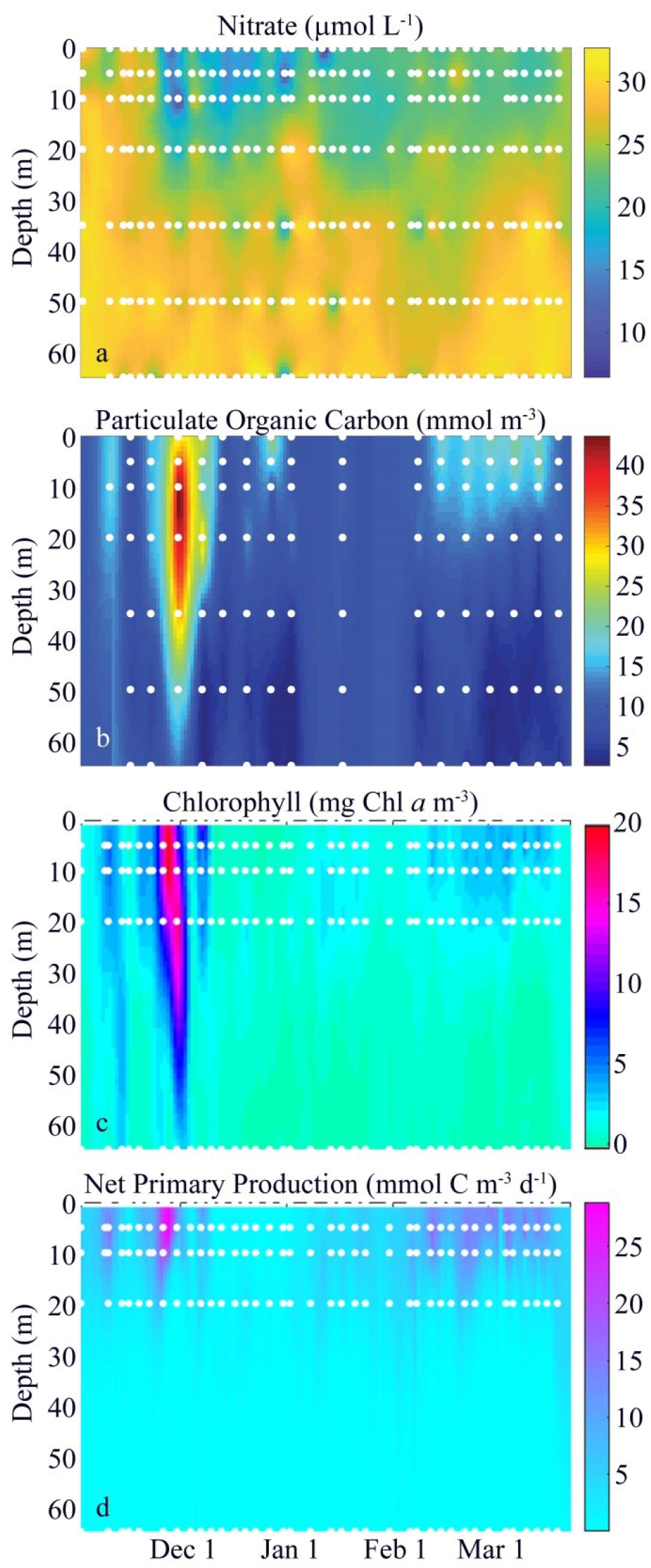
Chemical and biological properties at Station E during the Palmer 2012-2013 field season. a) nitrate, b) particulate organic carbon, c) Chl *a*, d) net primary production (corrected for mismatches between sampling depth and incubation light level, see methods). White dots are sampling points.

Throughout the season, sediment-trap derived carbon flux ranged from 5.8 ± 1.0 mmol C m^-3^ d^-1^ (mean ± standard deviation of triplicate samples) to 15.3 ± 1.5 mmol C m^-3^ d^-1^ (Fig. 4). Euphausiid fecal pellets were responsible for 56% of total carbon flux, although their contribution was highly variable and ranged from 2.5% to 128%. Similarly, >200-μm particles (which were primarily composed of fecal pellets, although pellets were also found in the <200-μm fraction) were responsible for 49.6% of total carbon flux and ranged from 4.1% to 65%. The above results should be interpreted with some caution, however, because simultaneous ^234^Th measurements suggested that the traps were under-collecting sinking particles by 29% (see below).

**Fig. 4. –.**
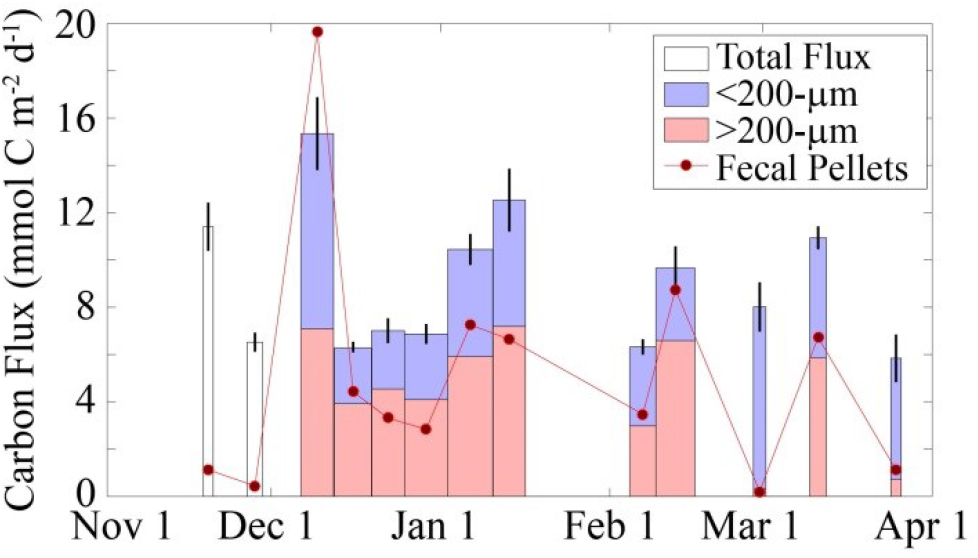
Sediment trap flux at 50 m depth. White, light blue, and light red boxes are total flux (non-size fractionated), flux of <200-μm particles, and >200-μm particles, respectively. Width of bars corresponds to deployment duration. Black lines are standard deviation of triplicate measurements. Red line and circles are fecal pellet mass flux in the sediment traps (no replicates).

### 3.2 Thorium dynamics and C:Th ratios

^234^Th activity was near equilibrium with ^238^U (2.4 dpm L^-1^) at the beginning of the field season and decreased substantially in the upper 20 m during the spring phytoplankton bloom (Fig. 5a, Supp. Table 1). Immediately following the bloom decay, ^234^Th concentrations increased briefly in January before remaining low in the upper 20 m until March.

**Table 1.** Parameters for the POC-FP models: first order thorium sorption coefficient (k_sorp,1_), second-order thorium sorption coefficient (k_sorp,2_), thorium desorption coefficient (k_desorp_), fraction of ingested thorium egested by zooplankton (Eg_Th_), and deviance information criterion (DIC).

|  | POC-FP<br>First Order | POC-FP<br>Second Order |
| --- | --- | --- |
| $k_{\text{sorp},1} \text{ (d}^{-1}\text{)}$ | $0.14 \pm 0.01$ | - |
| $k_{\text{sorp},2} \text{ (m}^3 \text{ mmol C}^{-1} \text{ d}^{-1}\text{)}$ | - | $0.0047 \pm 0.0002$ |
| $k_{\text{desorp}} \text{ (d}^{-1}\text{)}$ | $0.54 \pm 0.05$ | $0.0167 \pm 0.0076$ |
| $E_{\text{gTh}}$ | $0.68 \pm 0.02$ | $0.94 \pm 0.02$ |
| DIC | 1019 | 655 |

**Fig. 5. –.**
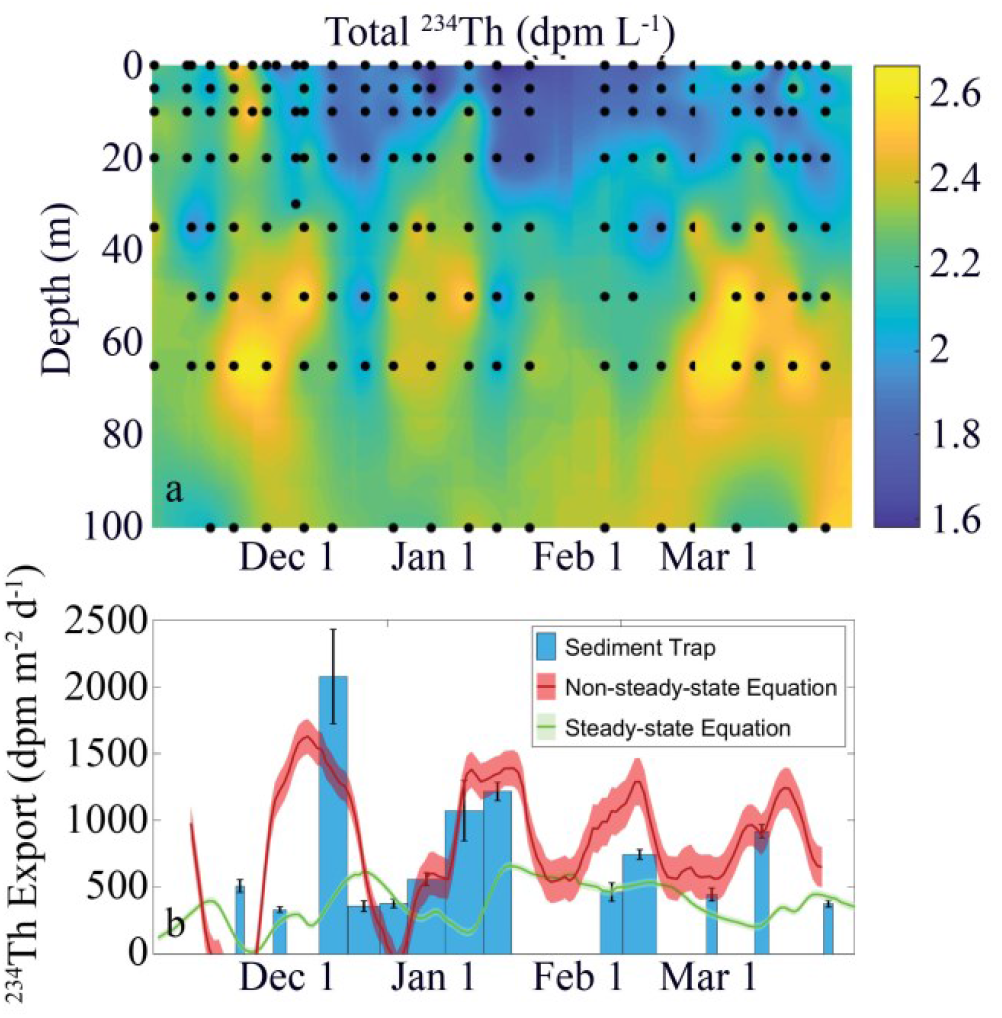
Total water column thorium activity (a) and export (b) during the 2012-2013 field season. In (b) blue bar plot shows direct measurements of ^234^Th flux measured in sediment traps deployed at 50-m, green line plot shows ^234^Th export estimated with a simple steady-state equation, and red line plot shows 2-week average ^234^Th flux estimated at 50-m depth from water column ^234^Th measurements using a non-steady state equation that accounts for vertical introduction of ^234^Th by diffusion.

Particulate ^234^Th activity seasonal patterns were largely driven by the spring bloom, during which time particulate ^234^Th activity reached >0.8 dpm L^-1^ and the percentage of total ^234^Th contained in particles reached ∼40%. During the rest of the season, particulate ^234^Th activities were ∼0.2 dpm L^-1^ and only ∼10% of total ^234^Th (Fig. 6, Supp. Table 2).

**Fig. 6. –.**
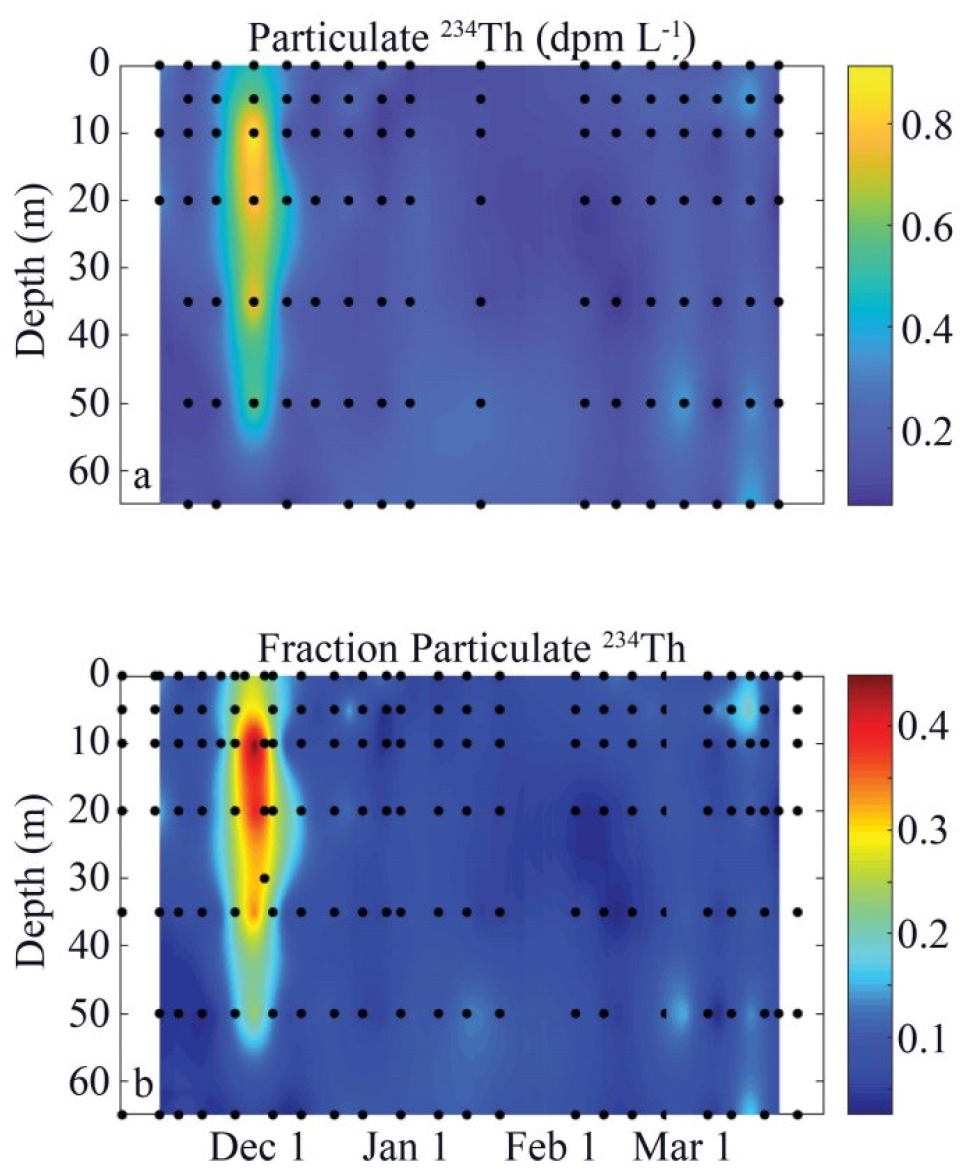
Particulate ^234^Th activity in the water column. b) ratio of particulate ^234^Th to total water column ^234^Th. Black dots show sampling points.

The C:^234^Th ratio of suspended particles was consistently higher in the upper euphotic zone (where it typically ranged from 40 – 80 μmol C dpm^-1^, but occasionally reached values greater than 100 μmol C dpm^-1^) than deeper in the water column (where values typically ranged from 5 – 40 μmol C dpm^-1^, Fig. 7a). C:^234^Th ratios exhibited different seasonal patterns in the upper and lower water column. Early in the season, C:^234^Th ratios were relatively constant with depth, but they increased slightly in the surface waters during the bloom and decreased substantially in deep waters beneath the bloom. C:^234^Th ratios in surface waters were slightly higher during the late summer bloom and particularly at the end of the field season than they were during the spring bloom or low biomass period in January and early February.

**Fig. 7. –.**
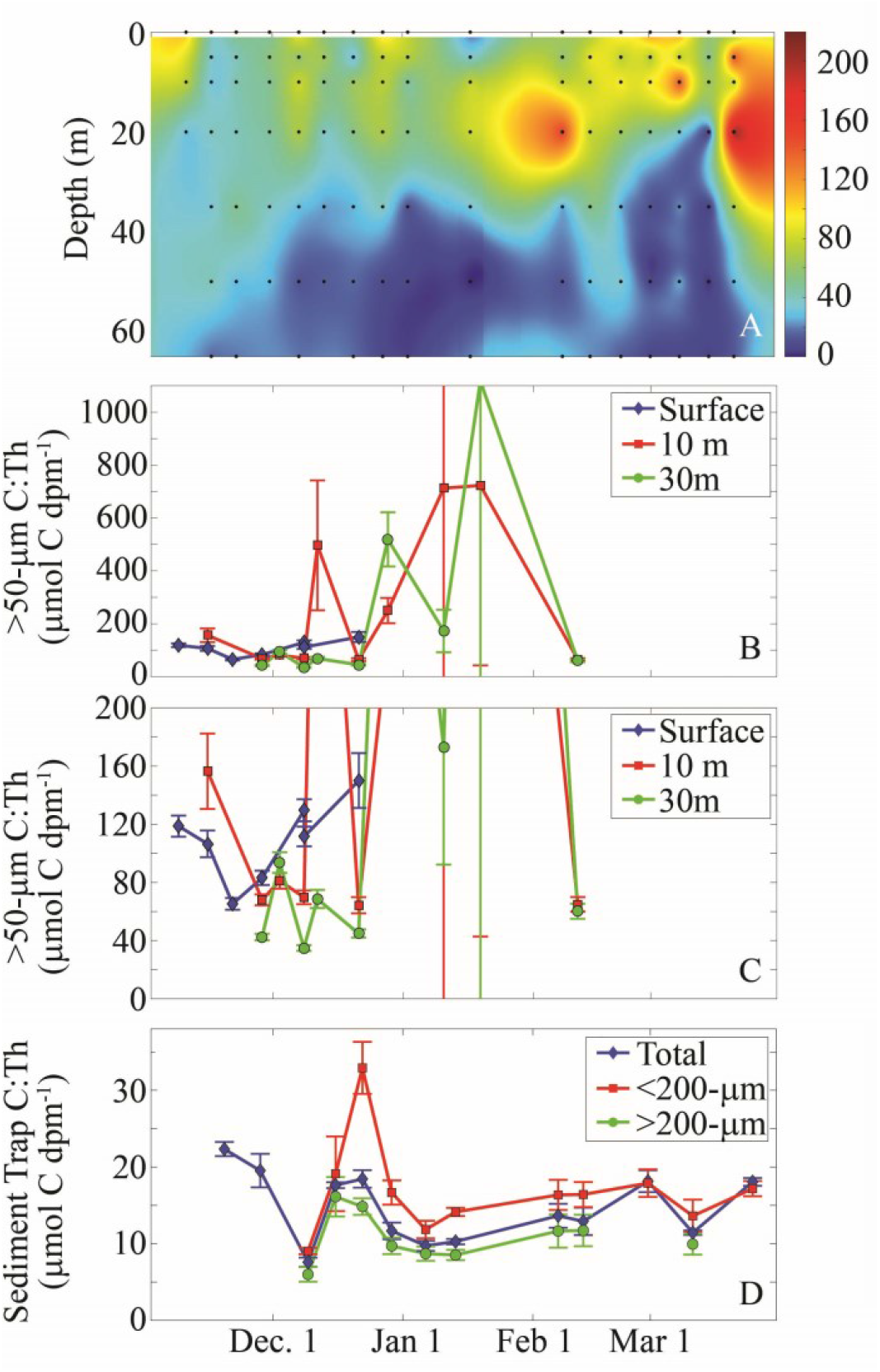
C:^234^Th ratios of: a) bulk particles in the water column, b) >50-μm size-fractionated particles collected through either surface sampling or using a Monsoon pump, c) same as (b) but with y-axis modified to highlight lower values, and d) sinking particles collected by sediment trap.

Size-fractionated (>50-μm) C:^234^Th ratios showed relatively low variability between the mixed layer and a depth of 30 m (Fig. 7b,c). However, they were almost always greater than the C:^234^Th ratios of bulk suspended particles. The C:^234^Th ratio of large (>50 um) particles ranged from ∼40 – 160 μmol C dpm^-1^ prior to and during the spring bloom. During the post-bloom low biomass period, however, the C:^234^Th ratios often exceeded 200 μmol C dpm^-1^, although there was often substantial uncertainty associated with these measurements, because ^234^Th activities were at times quite low despite a substantial amount of carbon on the filters. Although we did not undertake careful microscopic examination of the contents of the >50-μm samples, brief inspections (intended to ensure that no large metazoan zooplankton were present) did not notice the conspicuous euphausiid fecal pellets that often dominated sinking flux collected from the sediment traps. It thus seems likely that the >50-μm samples collected from the water column were qualitatively different from the dominant sinking particles.

In contrast to large or bulk suspended particles, the C:^234^Th ratio of sinking particles collected by the sediment trap at 50 m depth had low C:^234^Th ratios and comparatively low variability throughout the season (Fig. 7d, Supp. Table 3). The C:^234^Th ratio of sinking particles declined from 22.3 μmol C dpm^-1^ in the beginning of the field season to 7.6 μmol C dpm^-1^ near the end of the spring bloom. For the remainder of the summer it remained within a relatively narrow range from 9.7 to 18.4 μmol C dpm^-1^. Large (>200-μm) sinking particles (predominantly fecal pellets) consistently had a lower C:^234^Th ratio than smaller sinking particles in stark contrast to the pattern of generally higher C:^234^Th ratios found in >50-μm particles collected in the water column relative to bulk suspended particles. C:^234^Th ratios were also measured on three samples from which all non-fecal pellets had been removed. These fecal pellet samples had C:^234^Th ratios ranging from 10.9 to 11.8 μmol C dpm^-1^.

Direct comparison of ^234^Th fluxes into sediment traps to estimates of ^234^Th flux based on a non-steady state equation (Eq. 1) showed reasonably good agreement (Fig. 5b). In particular, both approaches determined a peak in export in early December, a subsequent decline in export after the decline of the spring bloom (late December to early January) and a subsequent increase in export in late January. The largest discrepancy between the two estimates occurred in mid-November (during the first sediment trap deployment), when the non-steady state equation actually predicted slightly negative ^234^Th sinking flux, because ^234^Th activity was increasing in the water column. Overall, point-to-point comparisons suggested that the sediment traps were on-average underestimating sinking flux by 29% relative to the non-steady state equation. This under-collection was likely related to the use of a moored configuration for the traps, because identical traps deployed in a surface-tethered configuration have been found to have no substantial over- or under-collection bias (Morrow *et al*., 2018). Notably, the use of a simple steady-state equation to estimate export (e.g., E = (^238^U-^234^Th)×λ_234_) substantially underestimates both total ^234^Th flux and its variability throughout the season (Fig. 5b, green line). Unsurprisingly, a steady-state model also lags true export flux because it estimates export averaged over the prior month.

### 3.3. Processes driving seasonal variability in C:^234^Th ratios

There was a strong negative correlation between the percentage of sinking organic carbon contributed by fecal pellets and the C:^234^Th ratio of sinking particles (Fig. 8). This is not surprising, since: 1) fecal pellets had lower C:^234^Th ratios than bulk suspended POM in the overlying water column, 2) large sinking particles (which were primarily fecal pellets) consistently had lower C:^234^Th ratios than smaller sinking particles (<200-μm) which contained a mixture of different particle types including diatoms, fecal pellets, and phytodetritus, and 3) fecal pellets were a dominant but highly variable contributor to export flux. The dominant role of fecal pellets also helps explain the lack of a clear seasonal trend in sediment trap C:^234^Th; *Euphausia superba* (the most abundant euphausiid in the WAP) is a highly mobile organism, with horizontal migrations and aggregating behavior that leads to high spatial and temporal patchiness. Nevertheless, it is surprising that the apparent increase in C:^234^Th ratios of suspended material in the surface layer was not reflected in the C:^234^Th ratio of sinking material and that the C:^234^Th ratio of sinking particles was so much lower than the C:^234^Th ratio of large particles collected from the water column.

**Fig. 8. –.**
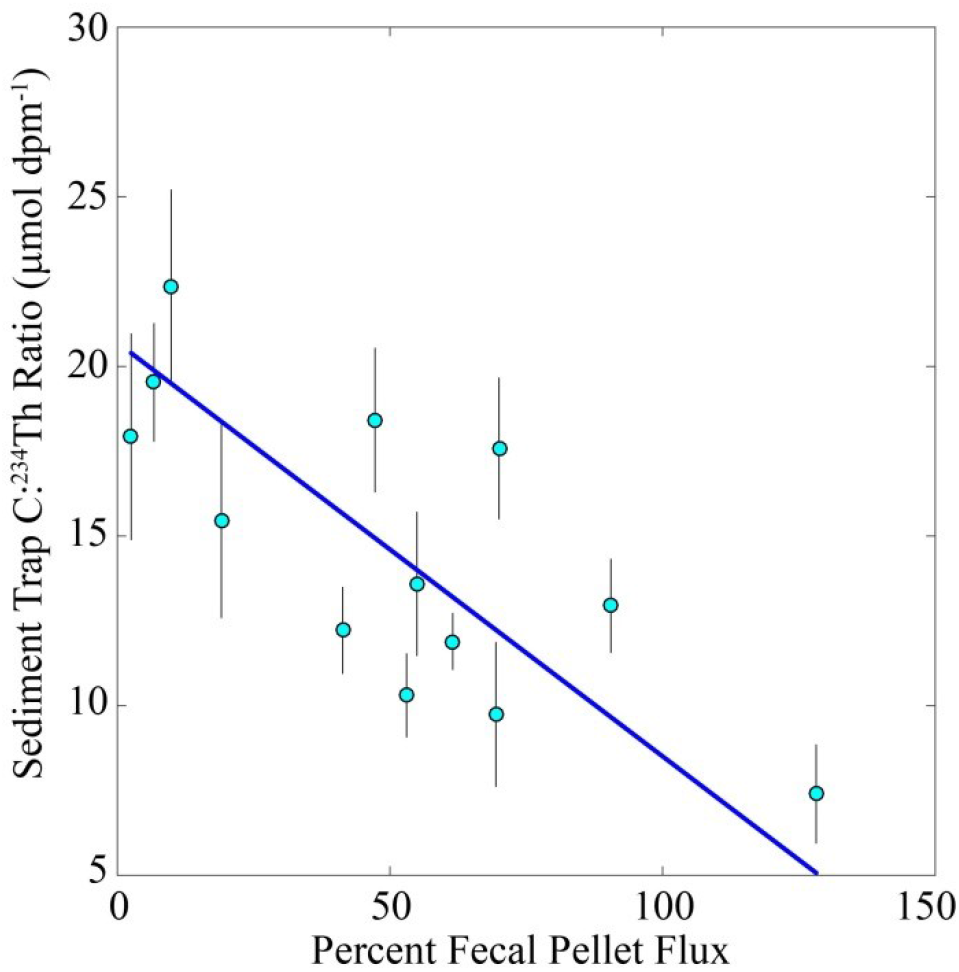
Relationship between the percentage contribution of fecal pellets to sediment trap carbon flux and the C:^234^Th ratio of particles collected in the sediment trap. Blue line is a Type II geometric mean regression: y = mx + b, where m = −0.12 ± 0.03 and b = 20.7 ± 1.6, r^2^ = 0.56.

The C:^234^Th ratio of suspended material was primarily controlled by POC and Chl *a* concentration (Fig. 9). The fraction of ^234^Th attached to particles was strongly correlated with POC (Pearson’s linear correlation, ρ = 0.75, p<<10^−5^). However, this relationship was only noticeable at comparatively high POC concentrations. When POC concentrations were low (<10 μmol C L^-1^), the fraction of ^234^Th bound to particles varied from low (∼3%) to moderate (∼15%) without a strong correlation to POC. At higher POC concentrations, as much as 45% of the ^234^Th was adsorbed onto particles. Chl *a* concentration was actually more strongly correlated with the fraction of ^234^Th attached to particles (Fig. 9a) with Pearson’s ρ = 0.83 (p<<10^−5^). The C:^234^Th ratio of suspended material was also influenced by both POC and Chl *a*. Particularly at low POC concentrations (<8 μmol C L^-1^) there was a strong positive correlation between C:^234^Th and POC (Fig. 9b). Across the full range of sampling points, however, the relationship was weaker (Pearson’s ρ = 0.48, p<<10^−5^), because at higher POC concentrations the C:^234^Th ratio was very sensitive to Chl *a* concentration. At similar POC concentrations, high Chl *a* corresponded to lower C:^234^Th ratios. This likely resulted from a greater proportion ^234^Th being adsorbed to particles when Chl *a* was higher.

**Fig. 9. –.**
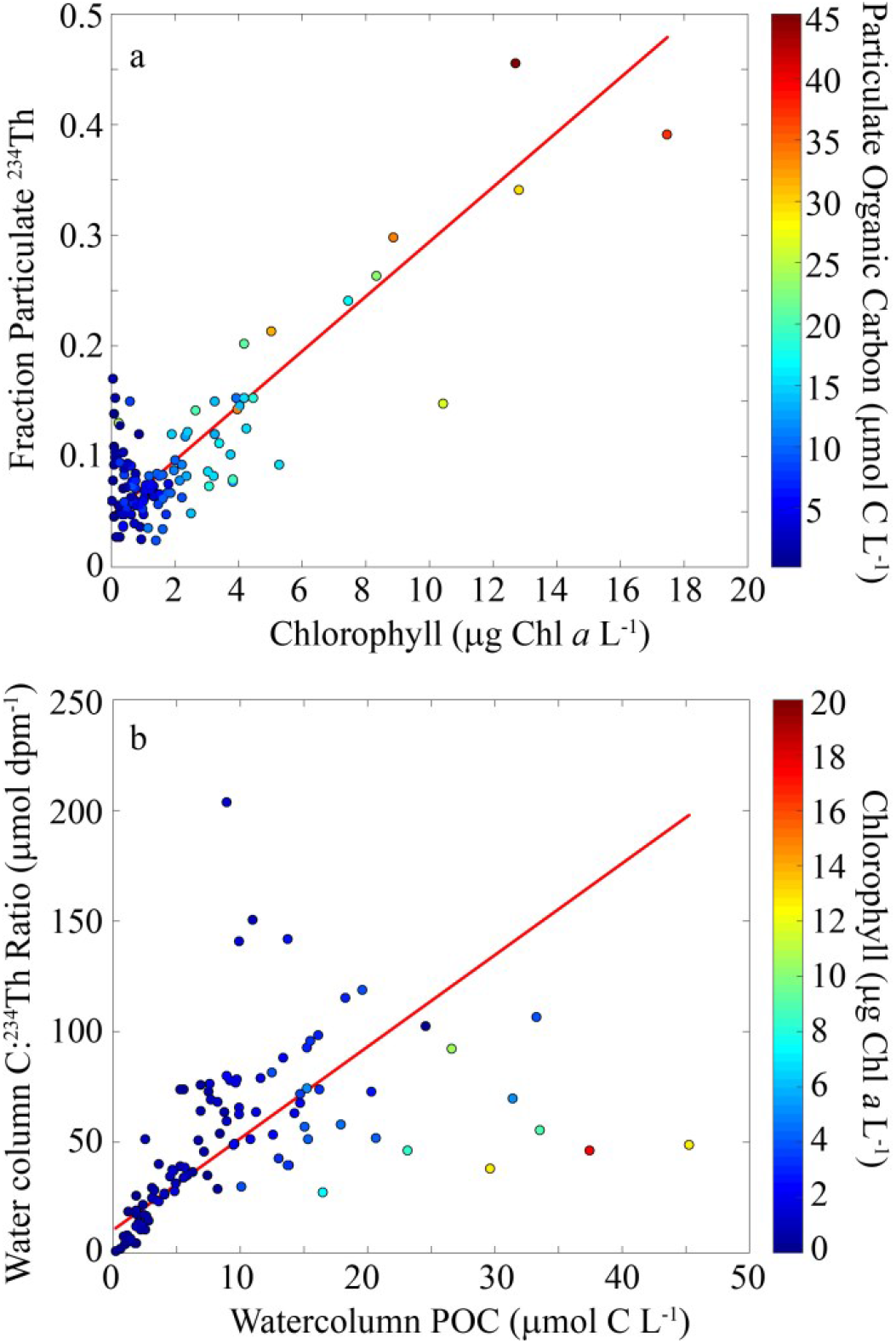
Particulate thorium relationships. a) Relationships between the fraction of ^234^Th bound to >1-μm particles and chlorophyll concentration. Color axis is POC (μmol C L^-1^). Red line is a Type II geometric mean linear regression, y = mx + b, where m = 0.025 ± 0.001 and b = 0.047 ± 0.005, r^2^ = 0.83. b) Relationship between the organic C:^234^Th ratio of particles in the water column and POC concentration. Color axis is Chl (mg Chl *a* m^-3^). Red line is a Type II geometric mean regression: y = mx + b, where m = 4.1 ± 0.4, b = 10.2 ± 5.3, r^2^ = 0.23.

### 3.4 Model-data comparisons and sorption kinetics

The Bayesian Markov Chain Monte Carlo parameter selection approach allowed us to fit thorium sorption models that reasonably simulated the observations. DIC was lowest for the POC-FP model with second order thorium sorption kinetics indicating that this model is best supported by the data (Supp. Table 4, Fig. 10). Notably, DIC was lower for all second-order thorium sorption kinetics models relative to the comparable first-order kinetics model, and these differences were often quite large; the only pair of models for which the DIC difference was less than 250 was for the POC model (for which DIC for the second-order kinetics model was 47 lower than for the first-order model). The POC-FP second-order kinetics model (and the second-order kinetics class of models generally) were able to capture key aspects of the thorium system including the high fraction of ^234^Th adsorbed to particles, the substantially lower C:^234^Th ratio of sinking particles collected in the sediment trap, and variability in the C:^234^Th ratio throughout the season (Fig. 11b,d,f,h). This model did, however, slightly overestimate the C:^234^Th ratio of suspended particles near the surface during the spring phytoplankton bloom. In contrast, the POC-FP first-order kinetics model (and the first-order kinetics class of models generally) did not accurately capture variability in the fraction of total ^234^Th bound to particles, but rather estimated that a relatively invariant ∼20% of ^234^Th was bound to particles at all depths and times (Fig. 11c). Consequently it substantially overestimated the C:^234^Th ratios of suspended and sinking particles during the bloom.

**Fig. 10. –.**
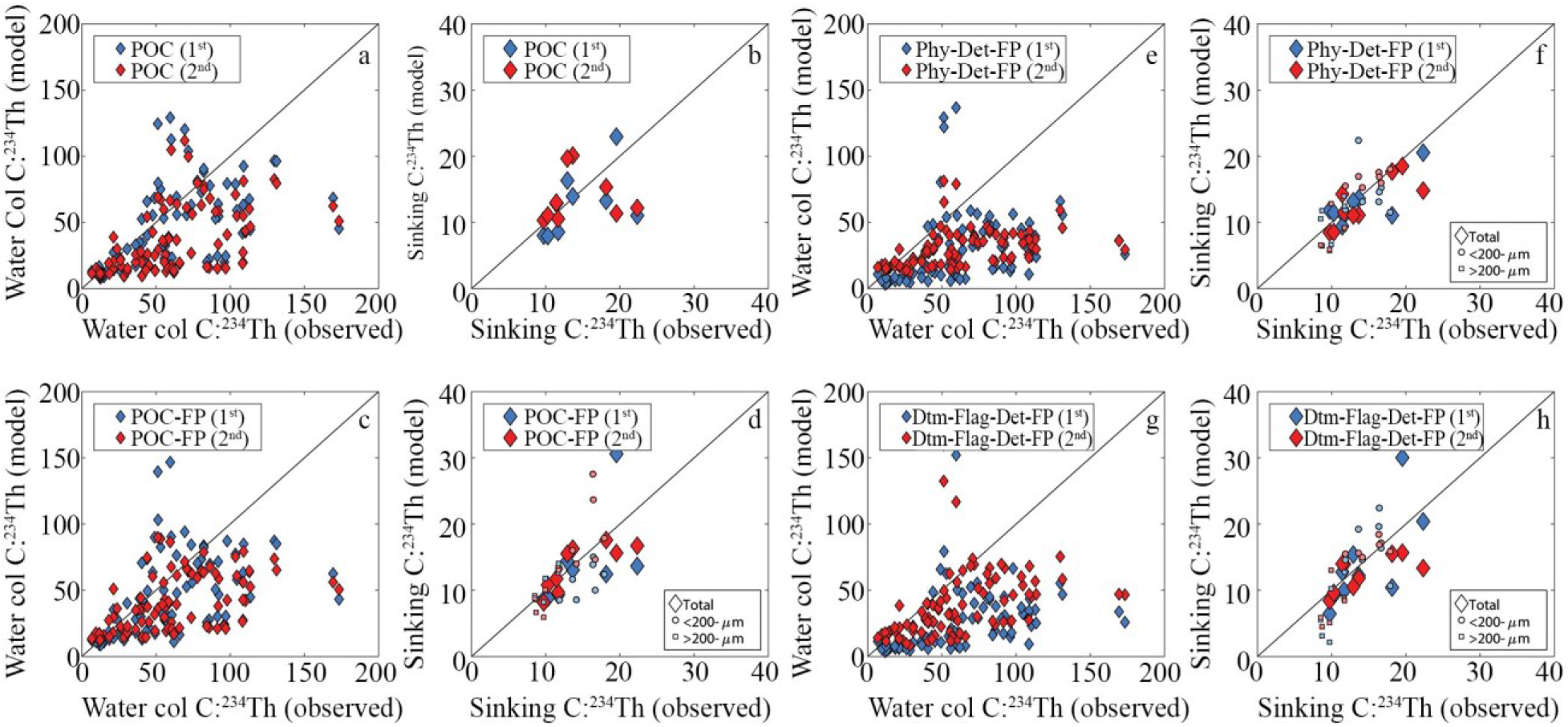
Model-observation comparisons for the POC models (a, b), POC-FP models (c,d), Phy-Det-FP models (e,f), and Dtm-Flag-Det-FP models (g,h). Panels a, c, e, and g show comparisons for suspended particles cfrom the water column. Panels b, d, f, and h show comparisons for sinking particles. In all panels blue symbols are for first-order kinetics model and red symbols are for second-order kinetics symbols.

**Fig. 11. –.**
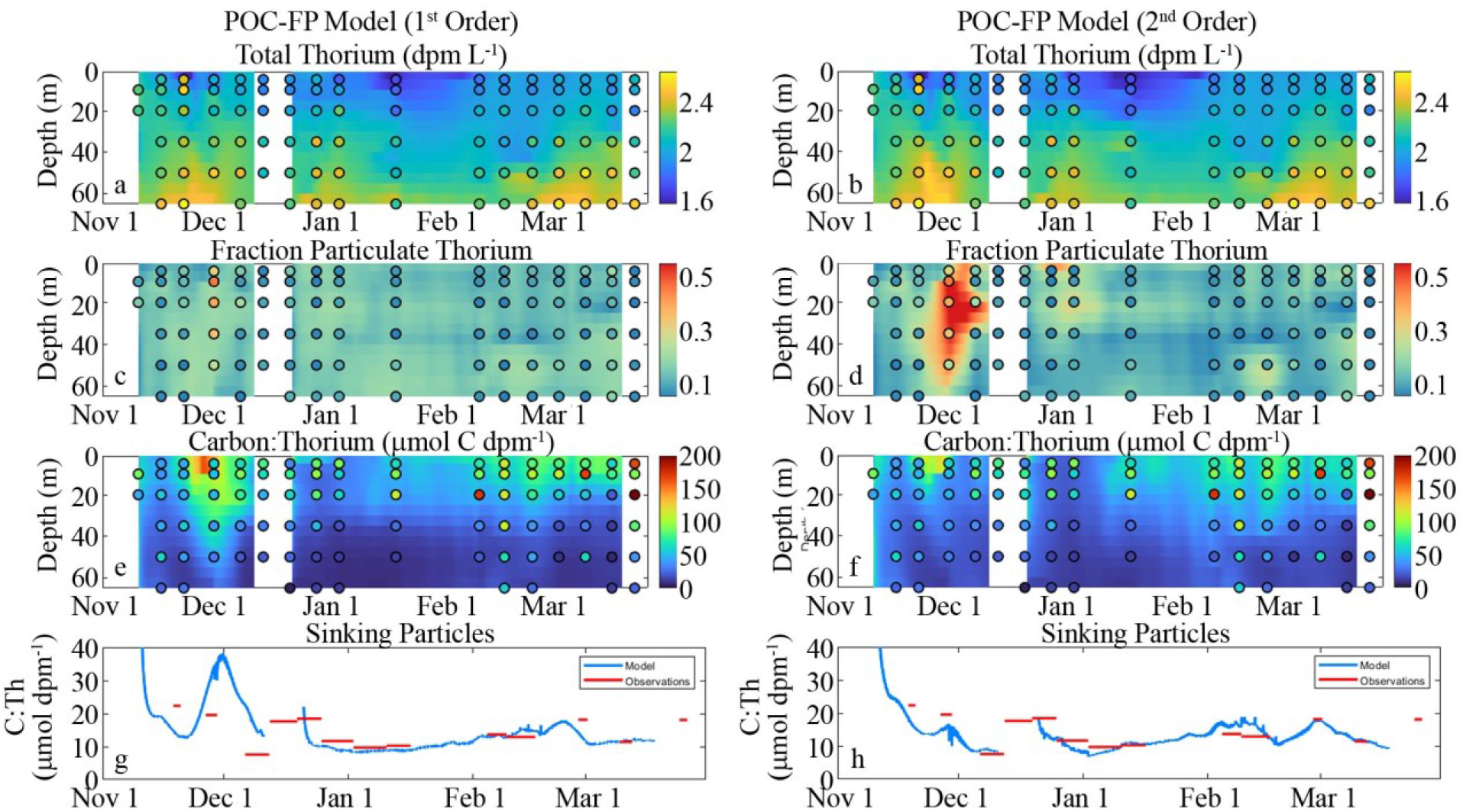
Model-observation comparisons for the POC-FP 1^st^ order model (a, c, e, g) and the POC-FP 2^nd^ order model (b, d, f, h). Smooth fields are model output. Black outlined circles are observations on the same color axis. Total (dissolved + particulate) ^234^Th (a, b). Fraction of total ^234^Th adsorbed to particles (c, d). C:^234^Th ratios of POC (e, f). C:^234^Th of sinking particles (g, h; blue lines = model; red lines = observations).

The models that included multiple particulate organic carbon compartments (Phy-Det-FP and Dtm-Flag-Det-FP) consistently underestimated the C:^234^Th ratios of total suspended particles, although they fairly accurately estimated the C:^234^Th ratios of sinking particles (Fig. 10). These results do not imply that the marine ecosystem behaves as a system with a single POC pool. POC is unquestionably a heterogeneous pool comprised of many living and non-living compartments spanning orders of magnitude differences in size and with many different surface properties likely leading to variable thorium sorption kinetics. Rather, our results suggest that, given our limited ability to constrain the true heterogeneity of this system, a simple model comprised of a single (mostly) suspended POC pool and a second class of more rapidly-sinking particles is a better predictor of C:^234^Th ratios than the more complex models that we tested.

### 3.5 Model parameters and processes affecting C:^234^Th ratios

Model results offer interesting insights to ^234^Th cycling in the WAP. We focus here on the second-order POC-FP model, which most accurately simulated the data. The model-fit thorium sorption parameter (k_sorp,2_) was 0.0047 ± 0.0002 m^3^ mmol C^-1^ d^-1^ (or 1.7 ± 0.1 m^3^ mmol C^-1^ y^-1^), which is about a factor of three smaller than our prior estimate for this parameter (Table 1). The estimated desorption parameter (k_desorp_) was 0.017 ± 0.0076 d^-1^ (or 6.1 ± 2.8 y^-1^), although we caution that this parameter was poorly constrained. Because ^234^Th decay is more rapid than desorption, the model was fairly insensitive to the desorption coefficient. The parameter that determines the fraction of thorium consumed by euphausiids that is egested as part of their fecal pellets (Eg_Th_) was 0.94 ± 0.02, indicating that most thorium consumed by euphausiids passes into their fecal pellets. For comparison, we assumed that only 30% of carbon consumed by euphausiids is egested (Eg_C_ = 0.3). This yields fecal pellets with a C:^234^Th ratio that is only about one third of the C:^234^Th ratio of euphausiid prey.

The model also offers insight into other dynamics of the thorium system. As mentioned, decay is a more important loss term for particulate ^234^Th than desorption. However, particle remineralization is actually the dominant loss term for particulate ^234^Th. Typical specific POC remineralization rates were in the range of 0.03 – 0.6 d^-1^ in the euphotic zone, compared to a decay constant of 0.028 d^-1^. Particle sinking was also an important loss term for particulate ^234^Th, with typical specific rates of loss from the euphotic zone of 0.01 to 0.02 d^-1^. Primary production also played an important role in shifting C:^234^Th ratios. Primary production drove typical specific POC production rates in the range of 0.1 to 1.0 d^-1^ in the euphotic zone. During periods of high primary production, the creation of new POC drove the C:^234^Th ratio higher than would be expected based on the steady state that would be anticipated if C:^234^Th were only affected by the processes of sorption, desorption, and decay.

Although the Dtm-Flag-Det-FP model was a meaningfully worse fit to the data than the POC-FP model, discussion of its parameterization and dynamics is still informative. The model predicted a substantially higher sorption coefficient for diatoms (0.015 ± 0.0005 d^-1^) than for flagellates (0.0037 ± 0.0008 d^-1^) or detritus (0.0038 ± 0.0003 d^-1^). Despite the higher sorption coefficient for diatoms, diatoms did not have a significantly lower C:^234^Th ratio than detritus. Both typically had C:^234^Th ratios in the range of 30 – 80 μmol C dpm^-1^. Conversely, detritus and flagellates had distinctly different C:^234^Th ratios despite similar sorption coefficients. Flagellate C:^234^Th ratios were often a factor of 5 greater than those for detritus. These similarities and differences between C:^234^Th ratios of phytoplankton and detritus were largely driven by the different formation processes and turnover times for these particles. Phytoplankton carbon is created through photosynthesis, which increases the C:^234^Th ratios of growing phytoplankton. Detritus, however, is formed from phytoplankton mortality and thus inherits the C:^234^Th ratio of existing phytoplankton, while continuing to adsorb more ^234^Th. Detritus also tends to have a longer residence time than phytoplankton, which allows it to reach a C:^234^Th ratio near the equilibrium that would be predicted from sorption, desorption, and decay processes.

## 4. DISCUSSION

The ^238^U-^234^Th disequilibrium approach has been widely used as a tool for investigating spatiotemporal variability in particle cycling at a range of scales including basin-scale (Owens et al., 2015; Puigcorbé et al., 2017), regional (Buesseler et al., 1995; Ducklow et al., 2018; van der Loeff et al., 2011), and mesoscale (Estapa *et al*., 2015; Resplandy *et al*., 2012; Stukel *et al*., 2017). Uncertainty in particle flux estimates associated with non-steady state dynamics and advective and/or diffusive transport of ^234^Th have been extensively studied (Buesseler et al., 1992; Ceballos-Romero et al., 2018; Dunne and Murray, 1999; Resplandy et al., 2012; Savoye et al., 2006), and general rules-of-thumb have been developed for identifying when such processes can be neglected and how uncertainties introduced by the use of simple steady-state, no-upwelling equations can be quantified. However, estimates of carbon flux from ^238^U-^234^Th disequilibrium are also complicated by variability in C:^234^Th ratios, which vary with depth, particle size and type, and sampling methodology, often over small spatial scales (Buesseler et al., 2006; Hung et al., 2012; Passow et al., 2006; Stukel et al., 2019). Unfortunately, the ship-time-intensity associated with measuring the C:^234^Th ratio of sinking particles often leads to much lower resolution sampling of the C:^234^Th ratio, relative to ^238^U-^234^Th disequilibrium. Many studies thus resort to applying a C:^234^Th ratio derived from measurements made at a single location and time to estimate carbon flux over a wide region (e.g., Ducklow et al., 2018; Estapa et al., 2015; Puigcorbé et al., 2017; Stukel et al., 2015). Without knowledge of the processes driving variability in C:^234^Th ratios, this introduces potentially large and poorly quantified uncertainty into estimates of carbon flux.

Clearly, empirical and/or mechanistic models that can predict changes in C:^234^Th ratios as a function of relevant biological and chemical parameters would greatly improve our measurements of the BCP. However, such approaches are complicated by the multitude of factors – many of which are not typically measured by the biogeochemists who study ^234^Th – that influence C:^234^Th ratios. Indeed, existing models used to study particle-thorium dynamics do not even agree about whether first-order kinetics (i.e., thorium scavenging rates are independent of particle concentration, Dunne et al., 1997; Lerner et al., 2016) or second-order kinetics (i.e., thorium scavenging rates are linearly dependent on particle concentration, Resplandy *et al*., 2012; Stukel and Kelly, 2019) are most appropriate to model thorium adsorption onto particles. Few attempts have been made to incorporate other information, such as particle size spectra, phytoplankton community composition or physiological status, or zooplankton dynamics (despite the presumed importance of these and other parameters), because of a paucity of studies that have quantified their impact. Our results provide new information about some of these processes.

The relative contribution of fecal pellets to total sinking flux was an important determinant of the C:^234^Th ratio of sinking particles. These fecal pellets had decreased (and relatively invariant) C:^234^Th ratios in comparison to the euphotic zone particles from which they were presumably formed. This makes sense in light of previous results that have shown that mesozooplankton typically have very high C:^234^Th ratios (Coale, 1990; Passow et al., 2006; Stukel et al., 2016; Stukel et al., 2019), and that the ^234^Th found in these organisms could bioaccumulate directly from dissolved ^234^Th (Rodriguez y Baena et al., 2008; Rodriguez y Baena et al., 2006). It thus seems likely that mesozooplankton preferentially assimilate carbon relative to ^234^Th, leaving their egesta enriched in ^234^Th. Based on the results of the Bayesian model parameterization analysis (with second-order-sorption-kinetics POC-FP model), we should expect the C:^234^Th ratios of fecal pellets to be only one third of the C:^234^Th ratio of the particles from which they were formed. The quantitative importance of euphausiid fecal pellets to total carbon flux in the Western Antarctic Peninsula found in this study and by Gleiber et al. (2012), thus leads to a dominant impact of these pellets on the C:^234^Th ratio of bulk sinking particles.

Nevertheless, the relative invariance of fecal pellet C:^234^Th ratios (Fig. 7d) remains surprising in light of the substantial variability in C:^234^Th ratios of both bulk and size-fractionated suspended particles (Figs. 7a-c). This may reflect variability in C:^234^Th ratios between different suspended particle classes in the euphotic zone. *Euphausia superba* (the dominant WAP euphausiid) feeds primarily on diatoms, which dominated the phytoplankton community (and POC) during the spring bloom in late Nov. to early Dec., but were comparatively scarce later in the season when the community was dominated by *Phaeocystis* and cryptophytes (Goldman *et al*., 2014; Kranz *et al*., 2015). The Dtm-Flag-Det-FP model suggested that diatoms had lower C:^234^Th than other phytoplankton (flagellates) and similar C:^234^Th ratios to suspended detritus. The increased C:^234^Th of bulk suspended particles later in the season, was in turn driven by an increase in the abundance of flagellates, which dominated the phytoplankton biomass in the summer and fall. It is possible that diatoms had lower C:^234^Th than either other phytoplankton or suspended detritus. This potentially explains the increased C:^234^Th ratios later in the season when diatoms were less abundant, as well as the comparatively decreased C:^234^Th ratios when Chl a concentrations are high seen in Fig. 9b. Mechanistically, the low C:^234^Th ratios of diatoms might be maintained through production of transparent exopolymers that contain many active binding sites for ^234^Th (Passow et al., 2006; Quigley et al., 2002; Santschi et al., 2003), leading to the strong correlation between Chl a and the fraction of ^234^Th adsorbed to particles seen in Fig. 9a.

Our results also offer strong support for the use of second-order rate kinetics models of thorium sorption. With first-order rate kinetics, we would expect the proportion of total ^234^Th that is bound to carbon to be relatively uncorrelated with POC concentration. This would lead to a proportional increase in C:^234^Th with increasing POC (or perhaps even a supralinear increase if total ^234^Th is lower during periods of high POC and export). However, Fig. 9b shows that a doubling of POC does not lead to a concomitant doubling of the C:^234^Th ratio. Furthermore, second-order rate kinetics models much more accurately estimated the fraction of ^234^Th bound to particles than first-order rate kinetics models (compare Fig. 6a to Figs. 10c and 10d), and the second-order rate kinetics POC-FP model had a DIC of only 653 (compared to 1012 for the first-order POC-FP model). Since differences of DIC on the order of 5 – 10 are typically considered meaningful, this offers exceedingly strong evidence that the second-order rate kinetics model more accurately simulates the system. The POC-FP (2^nd^ order) model suggests a thorium sorption coefficient of 0.0047 ± 0.0002 m^3^ mmol C^-1^ d^-1^. Previous estimates from a range of ecosystems spanning coastal and oceanic regions have ranged from 0.002 to 0.075 m^3^ mmol C^-1^ d^-1^, although most fall within a narrower range from 0.003 to 0.01 m^3^ mmol C^-1^ d^-1^ (Clegg *et al*., 1991; Clegg and Sarmiento, 1989; Clegg and Whitfield, 1993; Murnane *et al*., 1994; Stukel *et al*., 2019). These similar values across disparate regions suggest that second-order rate kinetics models are broadly applicable.

The striking difference between C:^234^Th ratios of >50-μm particles in the water column between the surface and 30 m depth and sinking particles collected by sediment traps at 50 m depth, is also an important result of this study. Although there was substantial uncertainty associated with C:^234^Th ratios of >50-μm particles measured in mid to late Jan. (as a result of very low ^234^Th content in these particles), C:^234^Th ratios of these particles were likely approximately an order of magnitude higher than the C:^234^Th ratio of sediment trap-collected particles. These varying results may derive from sediment traps and pumps sampling very different types of particles. Fecal pellets of *Euphausia superba* likely have sinking speeds that are hundreds of meters per day, suggesting that they spend minutes to hours in the euphotic zone before sinking out. In contrast, *Phaeocystis* colonies likely maintain their position in the euphotic zone, while many >50-μm aggregates may sink much more slowly than fecal pellets. These slowly-sinking (potentially high C:^234^Th) particle classes will thus be oversampled by pumps relative to their contribution to sinking flux. This highlights problems associated with inferring C:^234^Th ratios of sinking particles from measurements of size-fractionated particles. Indeed, McDonnell and Buesseler (2010) found that particle sinking speed in the WAP was essentially independent of particle size, while multiple studies have shown that <50-μm particles can contribute substantially to particle flux (Durkin et al., 2015; Hung et al., 2012). C:^234^Th ratios derived from size-fractionated in situ pump sampling should thus be interpreted with some caution.

## 5. CONCLUSIONS

C:^234^Th ratios of bulk suspended and size-fractionated particles varied throughout the spring-summer growing season at a coastal site near Palmer Station, Antarctica. The fraction of ^234^Th adsorbed onto particles was strongly correlated with both Chl *a* and POC concentrations. The correlation with Chl may imply an important role for diatoms (the dominant phytoplankton taxon during the spring bloom) in producing organic matter with active binding sites for thorium. C:^234^Th ratios of suspended particles generally increased in surface waters throughout the ice-free phytoplankton growing season and were correlated with POC concentration, particularly when POC concentrations were low. However, C:^234^Th ratios were lower than expected (based on high POC concentrations) during the height of the spring diatom bloom, as a result of strong scavenging of thorium onto particles at this time. The high variability in C:^234^Th ratios of suspended material was not reflected in the C:^234^Th ratios of sinking particles, which were relatively low and comparatively invariant throughout the spring, early summer, and late summer periods. C:^234^Th ratios of sinking particles were primarily driven by euphausiid fecal pellets, which were the dominant contributor of mass flux into sediment traps and had consistently low C:^234^Th ratios. These low C:^234^Th ratios likely result from preferential assimilation of carbon by mesozooplankton and hence elevated thorium activity in their egesta. A simple model including POC and fecal pellets that used second-order thorium sorption kinetics was able to simulate variability in C:^234^Th ratios of suspended and sinking particles throughout the season.

Our results highlight the importance of sinking particle composition in driving variability in the C:^234^Th ratio of sinking particles. The demonstrated impacts of diatoms and euphausiids (and potentially other taxa) on C:^234^Th ratios needs to be considered in studies that attempt to discern the functional responses of the biological carbon pump to plankton ecosystem dynamics. Studies that employ ^238^U-^234^Th disequilibrium to quantify carbon export over a large spatial domain commonly rely on less frequent measurement of the C:^234^Th ratio and hence extrapolate sparse C:^234^Th measurements across a heterogeneous ocean. Our results suggest that covariance between important particle-flux associated taxa and the C:^234^Th ratio will bias such studies. Higher spatial resolution sampling of the C:^234^Th ratio or more focused analyses of the mechanisms controlling the C:^234^Th ratio are thus crucial.

## ABBREVIATIONS

BCP: biological carbon pump
Chl: chlorophyll
dpm: decays per minute
POC: particulate organic carbon
WAP: Western Antarctic Peninsula
DIC: Deviance information criterion

## ACKNOWLEDGMENTS

We thank our many colleagues who participated in the Palmer Station 2012-2013 field season, especially Stef Strebel, Elizabeth Asher, Nicole Couto, Filipa Carvalho, Mikaela Provost, and Sven Kranz. This work was supported by NSF OPP awards 1340886 and 1440435 to HWD and 1951090 to MRS. HWD was also partly supported by a gift from the Vetlesen Foundation. Data used in this manuscript are available on the Palmer LTER Datazoo website (https://pal.lternet.edu/data) and in supplementary tables of this manuscript. Model code can be downloaded at: https://github.com/mstukel/Palmer_CTh_Model

## Notes

### Competing Interest Statement

The authors have declared no competing interest.

